# Kenyan Traditional Medicine: Exploring Old Solutions to the Modern Antibacterial Crises Through Natural Products Chemistry

**DOI:** 10.1101/2021.03.26.436821

**Authors:** Fidensio K. Ndegwa, Chaitanya Kondam, Debarati Ghose, Taiwo E. Esan, Zohra Sattar Waxali, Margaret E. Miller, Rangaswamy Meganathan, Nicholas K. Gikonyo, Paul K. Mbugua, Paul O. Okemo, Timothy J. Hagen

**Affiliations:** Department of Pharmacognosy, Pharmaceutical Chemistry and Pharmaceutical & Industrial Pharmacy, Kenyatta University, Nairobi, Kenya.; Department of Chemistry and Biochemistry, Northern Illinois University, DeKalb, IL, USA.; Department of Biological Sciences, Northern Illinois University, DeKalb, IL, USA.; Department of Plant sciences, Kenyatta University, Nairobi, Kenya.; Department of Microbiology, Kenyatta University, Nairobi, Kenya.

**Keywords:** Kenyan traditional medicine, antibacterial, infectious disease, extracts

## Abstract

Infectious diseases are a major cause of morbidity and mortality around the world, accounting for approximately 50% of all deaths in tropical countries. Despite remarkable progress in the field of microbiology, inability to control or mitigate, epidemics caused by drug-resistant microorganisms pose a serious health hazard to the global population. New therapeutic strategies must be developed as a global initiative for the prevention and control of infectious diseases. This study focuses on Kenyan medicinal plants and their activity against bacteria. Plant extracts obtained from seven Kenyan plants used in traditional medicine, were screened for their antibacterial activity against *Escherichia coli*, *Pseudomonas aeruginosa, Bacillus cereus, and Mycobacterium smegmatis*. Extracts from all these plants showed antibacterial activity against at least one of the tested organisms at a concentration of 2 mg/mL. Chemical screening showed the presence of different classes of phytochemicals such as alkaloids, terpenes, tannins, in some active extracts.

## 1. Introduction

The World Health Organization (WHO) reported infectious diseases are the major cause of illness and mortality around the world. (Manandhar *et al*., 2019) WHO estimated that about 50% of deaths in tropical countries are due to infectious diseases caused by bacteria (Mahady *et al*., 2005). The rate of mortality caused by multiple drug resistant (MDR), extensively drug resistant (XDR) and pan-drug resistant (PDR) microbes are increasing even in developed countries, such as the United States. Despite the remarkable progress in the field of microbiology, inability to control or mitigate, epidemics caused by drug-resistant microbes pose a serious public health hazard. The Centers for Disease Control (CDC) recommends various approaches for the control of infectious diseases such as vaccination, enhanced monitoring, diagnosis, and development of new therapeutics for the treatment of diseases. New therapeutic strategies must be developed as a global initiative for the prevention and control of infectious diseases (Manandhar *et al*., 2019).

For centuries, plants have been used for the treatment of various diseases and they can be investigated for the discovery of new antimicrobial agents. Many of the medicinal plant species are active against a wide range of microbes. In many developing countries such as Kenya, the use of medicinal plants for the treatment of various ailments is quite common. It is estimated that about 70% of the Kenyan population uses medicinal plants for primary health care. Divergent Kenyan communities like Ogiek, Taita, and Maasai use medicinal plants as therapeutics since they live far away from modern medical facilities. In the Ogiek community, 96 % of the population uses medicinal plants as their major therapeutic agents (Ndegwa, 2008). Various studies have reported the antimicrobial properties and efficacy of Kenyan medicinal plants (Nankaya *et al*., 2019). Evaluation of the medicinal properties of such plants can lead to the discovery and isolation of new therapeutic agents for the treatment of infectious diseases.

In this study, seven different Kenyan medicinal plants, *A. annua, A. remota, P. peruviana, P. africana, C. africana, S. didymobotrya*, and *B. micrantha* were chosen because these plants are commonly used for the treatment of malaria, pneumonia, diarrhea, and other diseases. Extracts were made from various plant parts using water, methanol, acetone, and hexane solvents and screened against four different strains of bacteria; *E. coli*, *P. aeruginosa, B. cereus* and *M. smegmatis* using the Kirby-Bauer disc diffusion assay (Goshu *et al.,* 2015).

## 2. Materials and Methods

The botanical information on the traditional medicinal plants of Kenya, used in this study are presented in Table1. Each plant sample was extracted with at least one or more of the following solvents: water, methanol, acetone and hexane. Depending on the availability, the samples were tested for antibacterial activity and the results are presented in Table 2. The antimicrobial compounds, kanamycin, nalidixic acid, and neem oil (*Azadirachta indica*) were used as positive controls. The phytochemical analysis and methods used in the preparation of plant extract samples are described in the supplemental material section.

**Table 1:**
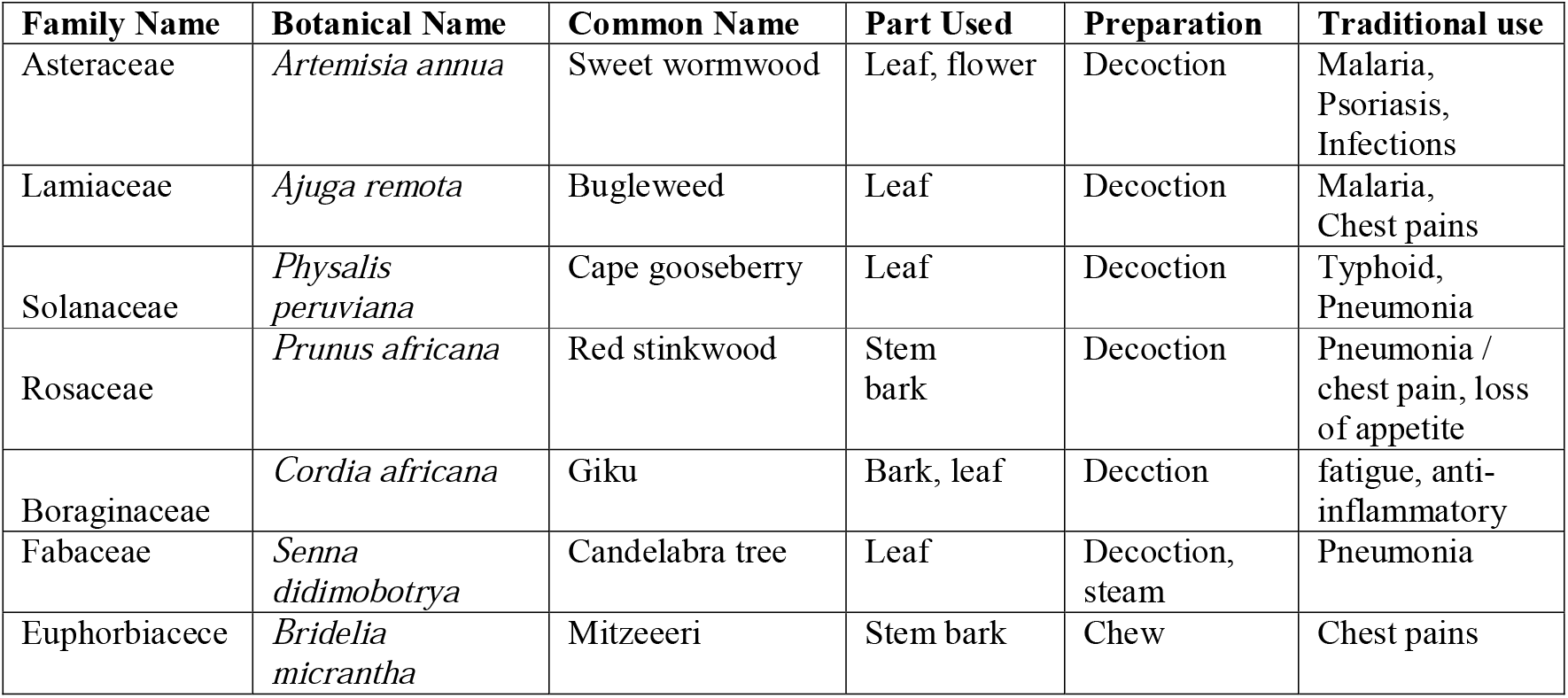
Medicinal plants tested for their antibacterial activity in the study. Kenyan medicinal plants, traditional preparation and use that were utilised in the study.

| Family Name | Botanical Name | Common Name | Part Used | Preparation | Traditional use |
| --- | --- | --- | --- | --- | --- |
| Asteraceae | *Artemisia annua* | Sweet wormwood | Leaf, flower | Decoction | Malaria, Psoriasis, Infections |
| Lamiaceae | *Ajuga remota* | Bugleweed | Leaf | Decoction | Malaria, Chest pains |
| Solanaceae | *Physalis peruviana* | Cape gooseberry | Leaf | Decoction | Typhoid, Pneumonia |
| Rosaceae | *Prunus africana* | Red stinkwood | Stem bark | Decoction | Pneumonia / chest pain, loss of appetite |
| Boraginaceae | *Cordia africana* | Giku | Bark, leaf | Decction | fatigue, anti-inflammatory |
| Fabaceae | *Senna didimobotrya* | Candelabra tree | Leaf | Decoction, steam | Pneumonia |
| Euphorbiaceae | *Bridelia micrantha* | Mitzeeeri | Stem bark | Chew | Chest pains |

**Table 2:** Determination of antibacterial activity. Diameter of zones of inhibition (ZOI) was (measured in mm.) for plant extracts against bacteria at 2 mg/mL and 0.2 mg/mL concentrations.

| Extract | *E. coli* |  | *P.aeruginosa* |  | *B. cereus* |  | *M.smegmatis* |  |
| --- | --- | --- | --- | --- | --- | --- | --- | --- |
| Dose in mg/mL | 2 | 0.2 | 2 | 0.2 | 2 | 0.2 | 2 | 0.2 |
| *A. annua* (water) | 11 | 0 | 11 | 0 | 10 | 0 | 11 | 0 |
| *A. annua* (acetone) | 10.5 | 0 | 19 | 0 | 18.5 | 7.5 | 0 | 0 |
| *A. annua* (hexane) | 8 | 0 | 0 | 0 | 15 | 0 | 12 | 0 |
| *A. remota* (water) | 0 | 0 | 8 | 0 | 7 | 0 | 0 | 0 |
| *B. micrantha* (water) | 10 | 0 | 10.5 | 0 | 16 | 0 | 16 | 0 |
| *B. micrantha* (methanol) | 11 | 0 | 14 | 0 | 17.5 | 7.5 | 11.5 | 0 |
| *B. micrantha* (acetone) | 0 | 0 | 12 | 0 | 16 | 0 | 15 | 0 |
| *B. micrantha* (hexane) | 8.5 | 0 | 12.5 | 0 | 15 | 0 | 19 | 0 |
| *C. africana* (water) | 0 | 0 | 9.5 | 0 | 7.5 | 0 | 9 | 0 |
| *C. africana* (methanol) | 0 | 0 | 13 | 0 | 10 | 0 | 12 | 0 |
| *C. africana* (hexane) | 10 | 0 | 0 | 0 | 17.5 | 7 | 9 | 0 |
| *P. peruviana* (water) | 0 | 0 | 12 | 0 | 7 | 0 | 11.5 | 0 |
| *P. africanus* (water) | 0 | 0 | 0 | 0 | 17 | 0 | 12.5 | 0 |
| *P. africanus* (acetone) | 10 | 0 | 10.5 | 0 | 20 | 0 | 0 | 0 |
| *P. africanus* (hexane) | 0 | 0 | 12.5 | 0 | 0 | 0 | 12 | 0 |
| *S. didymobotrya* (water) | 8.5 | 0 | 11.5 | 0 | 19.5 | 9 | 11 | 0 |
| *S.didymobotrya* (methanol) | 0 | 0 | 0 | 0 | 19 | 8.5 | 12 | 0 |
| *S. didymobotrya* (hexane) | 0 | 0 | 0 | 0 | 11 | 0 | 11 | 0 |
| Kanamycin (control) | 23 | 13 | 0 | 0 | 27 | 20 | 27 | 18 |
| Nalidixic acid (control) | 10 | 0 | 10 | 0 | 33 | 25 | 9 | 0 |
| *Azadirachta indica*<br>(Neem oil) (control) | 13 | 0 | 12 | 0 | 10 | 0 | 12 | 0 |

## 3. Results and discussion

It was observed that extracts of *Artemisia annua, Ajuga remota, Physalis peruviana, Prunus africanus, Cordia africana, Senna didymobotrya*, and *Bridelia micrantha* significantly inhibited growth of various bacteria as determined by Kirby Bauer disc diffusion method (Table 2).

*A. annua* extracts prepared with water were effective against all four organisms at 2.0 mg/ml. The acetone extract was twice as effective as the water extract against *P. aeruginosa* and *B. cereus*. *B. cereus* was very sensitive to acetone extract since it was inhibited even at 0.2 mg/ml. The hexane extract was ineffective against *P. aruginosa*.

Extracts of *B. micrantha* prepared with water, methanol and hexane were highly effective against all the four organisms. *B. cereus* was highly sensitive to methanol extract since it was inhibited even by the low concentration of 0.2 mg/ml. Surprisingly, *E. coli* was resistant to acetone extract in contrast to the other three organisms.

The *C. africana* extracts in water or methanol were active against *P. aruginosa*, *B. cereus* and *M. smegmatis* while *E. coli* was resistant. In contrast, the hexane extract was active against all organisms except *P. aruginosa. B. cereus* was very sensitive to the hexane extract since it was effective even at the low concentration of 0.2 mg/ml. The pattern of inhibition exhibited by the water extract of *P. peruviana* is similar to that of *C. Africana* methanol extract in that they both inhibited *P. aruginosa, B. cereus* and *M. smegmatis.*

Surprisingly, depending on the solvent used *P. africanus* extracts showed wide variation in their effect on the bacteria tested. The water extract inhibited only the two Gram positive *B. cereus* and *M. smegmatis. P. aruginosa, E. coli* and *B. cereus* were found to be sensitive to acetone extract in contrast to *M. smegmatis* which was resistant. The sensitivity of Gram-negative *P. aruginosa* and Gram-positive *M. smegmatis* for the hexane extract was about the same as shown by the diameter of the zone of inhibition.

The water extract of *S. didymobotrya* inhibited all the four organisms. In contrast, the hexane and methanol extracts were effective against the two Gram-positive bacteria tested. The methanol extract was very effective against *B. cereus* since it was inhibitory even at the low concentration of 0.2 mg/ml.

Phytochemicals are vital components of plants that are responsible for antifungal and antibacterial properties. *A. annua* extract showed anti-bacterial and anti-fungal activity (Rolta *et al*., 2021). Phytochemical screening showed the presence of tannins, steroids, carbohydrates, proteins/amino acids, flavonoids, anthraquinones, and terpenoids. *A. annua* plant extract also has antioxidant activity due to the presence of tannins (Bora *et al*., 2011).

Preliminary phytochemical screening of *A. remota* extracts showed the presence of tannins, flavonoids, steroids, carbohydrates, proteins, cardiac glycosides, anthraquinones, and terpenoids in the extracts. The extracts of *A. remota* were reported to show significant anti-diabetic effect as these phytochemicals enhance the activity of glycolytic enzymes (Tafesse *et al*., 2017). The extracts were also reported to have antidiarrheal and antimalarial activity (Yacob *et al.,* 2016). The steroidal metabolite ergosterol-5,8 endoperoxide isolated from this plant acts as a potential anti-bacterial agent (Yong, *et al.,* 2014).

*B. micrantha* extract contains, alkaloids, tannins, anthraquinones, steroids and flavonoids and were reported to exhibit antibacterial activity against various bacteria and reported to exhibit antioxidant and antihaemolytic properties (Adefuye *et al.,* 2013).

Chemical analysis *of C. africana* extracts showed the presence of flavonoids, tannins, steroids, carbohydrates, and cardiac glycosides. The extracts exhibited antibacterial, anti-inflammatory and antioxidant activity. Further, it was reported to exhibit antiulcer activity in pyloric ligated rats (Yismaw *et al.,* 2020).

*P. peruviana* contains tannins, steroids, anthraquinones, flavonoids, terpenoids by phytochemical analysis and reported to have anti-bacterial and anti-fungal activity. Extracts from various parts of this plant exhibited anti-parasitic effect against the malarial parasite activity against *Plasmodium falciparum*. (Kamau *et al*., 2020).

The stem bark of *P. africanus* has been used as a remedy for stomach-ache, as a laxative for cattle, and prostate cancer (Onyancha *et al;* 2018). Tannins and anthraquinones in the bark extract show antioxidant activity (Mwangi *et al*., 2018). Phytochemical analysis of the bark showed the presence of carbohydrates, tannins, flavonoids and terpenes. Stem bark extract showed an anti-inflammatory effect in mice and can be used to isolate potent anti-inflammatory compounds (Ngeranwa *et al.*, 2020).

Our phytochemical analysis revealed that *S. didymibotrya* contains steroids, alkaloids, tannins, carbohydrates, cardiac glycosides, anthraquinones, terpenoids and flavonoids. The leaf extracts were reported to show analgesic activity in mice models and reported to have anti-bacterial and anti-fungal activity (Jeruto *et al*., 2017).

## 3. Conclusion

In summary, the results presented in this report provide strong evidence that extracts of *A. annua, A. remota, P. peruviana, P. africanus, C. africana, S. didymobotrya*, and *B. micrantha* have antibacterial activity against *E. coli, P. aeruginosa, B. cereus and M. smegmatis.* Of particular significance is the effectiveness of extracts of every plant against *P. aeruginosa,* which is resistant to several antibiotics and an opportunistic pathogen of immunocompromised individuals and chronic lung diseases patients.

A significant finding is the effectiveness of the plant extracts against the Gram-positive pathogen, *B. cereus*. This bacterium is a spore former and usually causes food poisoning and eye infections. However, it is increasingly reported to cause fatal non-intestinal infections such as fulminant sepsis, central nervous system infections and hospital acquired infections, particularly in immunosuppressed patients. Examination of the results in Table 2 reveals that every plant examined, irrespective of the solvent used was inhibitory to *B. cereus* except for *P. africanus* hexane extract. Furthermore, four of the plant extracts from different solvents were inhibitory at 0.2 mg/ml. Many of the same plant extracts also inhibited the other Gram-positive bacterium *M*. *smegmatis* except for water extracts of *A. annua* and *A. remota* and acetone extract of *P. africanus.*

The very effective inhibitory properties of these plant extracts hold great promise in discovering new antibiotics. From the results in Table 2, it is obvious that these plant extracts were far more effective against the two Gram positive *B. cereus* and *M. smegmatis* than the two Gram-negative bacteria. These plant extracts can be used as prospective sources for the development of anti-microbial agents. Further attempts to identify and purify the active components from these extracts will give rise to promising antibacterial lead compounds that may be developed to meet the urgent need for new antibiotics.

## Supplemental material

Experimental details related to this paper is available online

## Acknowledgement

The authors acknowledge Northern Illinois University and Kenyatta University for supporting and funding this work.

