## Supplemental material for "Kenyan Traditional Medicine: Exploring Old Solutions to the Modern Antibacterial Crises Through Natural Products Chemistry"

**Experimental**

1. **Preparation of Plant material**

***Collection of plant material***

Dry powdered material of plants *Ajuga remota* , *Bridelia micrantha*, *Senna didymobotrya , Cordia* *africana* , *Physalis peruviana* , *Prunus africanus,* and *Artemisia annua* were obtained from the Department of Pharmacognosy, Pharmaceutical Chemistry and Pharmaceutical & Industrial Pharmacy, Kenyatta University, Nairobi, Kenya

***Preparation of extracts***

Dry powdered plant material was sequentially extracted using three solvents: water, methanol, and hexane. The dry powdered plant material was first extracted with water by decoction and then sequentially extracted with methanol and hexane in a Soxhlet apparatus. Plant material was also extracted non-sequentially using acetone with Soxhlet extraction. Each plant sample was extracted in at least one or more of the following solvents – water, methanol, acetone, and hexane. Then solvents methanol, acetone, and hexane evaporated under Nitrogen (N_2_) gas, and water were evaporated by lyophilization. Depending on sample availability and extraction yield, the samples were tested for antibacterial activity.

2. **Evaluation of antimicrobial activity**

***Preparation of discs***

The plant extract stock solutions were prepared with dimethyl sulfoxide (DMSO) (0.2 mg/mL, 2 mg/mL). Stock solutions of plant extracts with different concentrations permeated on 6 mm sterile paper discs (Becton Dickson, Australia). Paper discs permeated with DMSO stock solutions were used for testing activity.

***Microbial strains***

Strains of *Pseudomonas aeruginosa* - Gram-negative ATCC 27853, *Escherichia coli* Gram-negative ATCC 23846, *Mycobacterium smegmatis* Gram-positive NRRL-B-24020, *Bacillus cereus* Gram-positive Departmental stock were obtained from Department of Biology Northern Illinois University and stored in glycerated Luria-Bertani (LB) broth medium at -75° C until use.

***Preparation of inoculum***

Bacterial strains were incubated at either 37°C or 28°C depending on the optimum growth temperature of each bacterium. For preparing inoculum for the assay, cells were inoculated from an agar plate into 5ml. of sterile LB broth and incubated under shaking conditions overnight. Fresh overnight culture was then aseptically spread on LB agar plates using sterile cotton swabs.

***Kirby Bauer disc diffusion method***

Sterile filter paper disks were placed on the plates with forceps. Plant extracts were dissolved dimethyl sulfoxide (DMSO) and added at different concentrations on each disc paper. Kirby-Bauer disc diffusion susceptibility test was conducted for each plant extract at 0.2 mg/mL, and 2 mg/mL against *Pseudomonas aeruginosa* (ATCC 27853), *Escherichia coli* (ATCC 23846), *Mycobacterium smegmatis* (NRRL-B-24020), *Bacillus cereus* (NIU Departmental stock). The bacteria were grown in LB broth and spread on LB agar plates with cotton swabs. Sterile filter paper discs were placed on the plates with forceps. The plates were then incubated at 37°C for 24-48 hours depending on the organism and the diameter of the zone of inhibition was measured. For controls, in addition to traditional antibiotics kanamycin and nalidixic acid, an established plant-based extract that is extensively used as a pest and disease control agent called *Azadirachta indica* (commonly referred to as Neem oil) was also used.

3. **Phytochemical analysis**

Extractions were performed in a Soxhlet apparatus. Thin-layer chromatography employed silica gel plates F 1500/LS from Merck. Samples were spotted on the plates about 5µL using capillary tubes. Chromatographic spots were visualized using ultraviolet lamp emitting at 254 and 365 nm. All solvents and reagents were of analytical reagent grade.

***Qualitative Phytochemical Characterization***

Anisaldehyde/ Sulphuric acid reagent for Steroids (Touchstone, 1992); Dragendorff reagent for alkaloids (Touchstone, 1992); potassium hydroxide (Bornträger reaction) for coumarins (UV 365 nm) and anthraquinones (vis. and UV 365nm); 5% ethanolic solution of H_2_SO_4_ for cardiac glycosides (vis. and UV 365nm) (Starke et. al., 1998); Aluminum chloride solution (1% ethanolic AlCl_3_) for flavonoids (UV 365 nm) (Gage et. al., 1951); Iron (III) chloride reagent (3% FeCl_3_) for tannins and phenolic compounds (Fink et. al., 1959); Ninhydrin for amino acids, amines, and amino sugars (0.2% ethanolic ninhydrin solution) (Famy et. al., 1961); Phenol / sulfuric acid solution for carbohydrates (Touchstone, 2019); Vanillin / H_2_SO_4_ solution for Terpenes/Terpenoids (Jiang et. al., 2016). The qualitative phytochemical characterization and analysis data is shown in Table 1.

***Thin-layer chromatography***

The volume of the spots applied on the chromatographic plates was about 5µL. Then TLC plates were treated with the reagents for colorimetric detection of phytochemicals.

**Table 1: Phytochemical analysis of plant extracts**

| Plant Extract | Steroids | Alkaloids | Tannins  /phenols | Carbohydrates | Proteins  /amino acids | Cardiac lycosides | Flavonoids | Anthraquinones | Terpenes/Terpenoid |
| --- | --- | --- | --- | --- | --- | --- | --- | --- | --- |
| ***Ajuga remota*** | | | | | | | | | |
| Water | + + + | - | + | + + + | + + + | + + + | + + + | + + + | + + + |
| Methanol | + + | - | + | + + | + | + + | + + | + + | + + |
| Acetone | + + | - | - | - | - | + | - | + | + + |
| Hexane | + | - | - | - | - | - | - | - | + + |
| ***Artemisia annua*** | | | | | | | | | |
| Methanol | + + | - | + + | + | + | + | + + | + + + | + + |
| Acetone | + + + | - | + + | + | - | + | + | + + + | + + |
| Hexane | + | - | - | - | - | - | - | - | + + |
| ***Prunus africana*** | | | | | | | | | |
| Methanol | + | - | + | + | - | + | + | + | + |
| Acetone | + + + | - | + + | + | - | + + + | + | + + + | + |
| Hexane | - | - | - | - | - | - | - | - | + |
| ***Physalis peruviana*** | | | | | | | | | |
| Acetone | + + + | - | + | + + | + | + | + + | + | + + + |
| Hexane | + | - | - | - | - | - | - | - | + + + |
| ***Cordia africana*** | | | | | | | | | |
| Water | + + | - | + + | + + | + + | + | - | + | - |
| Methanol | + + + | - | - | + | - | + | - | - | - |
| Hexane | + | - | - | - | - | + | - | - | + |
| ***Senna didymibotrya*** | | | | | | | | | |
| Water | + + + | - | + | + + + | + | + + + | + | + | - |
| Methanol | + + | + | + + | + + | - | + + | + + | + + + | - |
| Acetone | + + + | - | - | - | - | + | + | + | - |
| Hexane | - | - | - | - | - | - | - | - | + |
| ***Bridelia micrantha*** | | | | | | | | | |
| Water | + + | + | + + + | + | + | - | - | + | - |
| Methanol | + + + | + + | + + + | + + | + | + + | - | + + + | - |
| Acetone | + + + | - | + + | - | - | - | - | + | - |
| Hexane | - | - | - | - | - | - | - | - | + |

**+ = low presence ++ = moderately present, +++ = highly present, - = absent**
